# Psoralen Enhances Direct Cardiac Reprogramming via PPARα Activation and Promotion of Mitochondrial Fission

**DOI:** 10.1101/2025.10.08.681288

**Authors:** Wenjie Li, Haixia Liu, Xinyu Wan, Ding Cheng, Zhiguo Zhang, Ruyuan Zhu

**Affiliations:** Institute of Basic Theory for Chinese Medicine, China Academy of Chinese Medical Sciences, Beijing 100700, China

**Keywords:** Psoralen, Direct cardiac reprogramming, PPAR α, UCP1, Mitochondrial fission

## Abstract

**BACKGROUND:** Cell-based therapy is a promising strategy for heart repair and regeneration. However, its clinical application has been hampered by the low efficiency of cell direct reprogramming and the immature state of the inducing cells. Developing novel strategies to enhance direct reprogramming efficiency and yield mature functional cardiomyocytes remains a critical challenge.

**METHODS:** We evaluated the effect of Psr on cardiac reprogramming using RepSox and Forskolin (RF) as the baseline cocktail. Reprogramming efficiency, cardiomyocyte marker expression, and calcium handling were assessed by RT-qPCR, immunofluorescence. Ultrastructural features were examined by transmission electron microscopy. Metabolic profiles were analyzed using Seahorse assays. Transcriptomic changes were explored by RNA sequencing, followed by pathway and protein–protein interaction analyses. A myocardial infarction (MI) mouse model was used for in vivo validation.

**RESULTS:** Supplementation with Psr (10 μM) markedly enhanced the induction of induced cardiomyocytes (iCMs), leading to earlier appearance of beating clusters (day 1 vs. day 6–8), improved sarcomere organization, robust calcium transients, and higher energy metabolism. Transcriptomic profiling identified activation of the PPAR signaling pathway, with PPARα, RXRG, and UCP1 as central regulators. Mechanistically, Psr promoted mitochondrial fission, thereby facilitating metabolic remodeling essential for cardiomyocyte maturation. In vivo, RF+Psr treatment significantly improved cardiac function and reduced fibrosis after MI compared to RF alone.

**CONCLUSION:** Psr enhances direct cardiac reprogramming by activating PPAR signaling and promoting mitochondrial fission. These findings provide a novel mechanistic framework and suggest psoralen as a promising natural enhancer for cardiac regeneration strategies.

**Graphical abstract:** 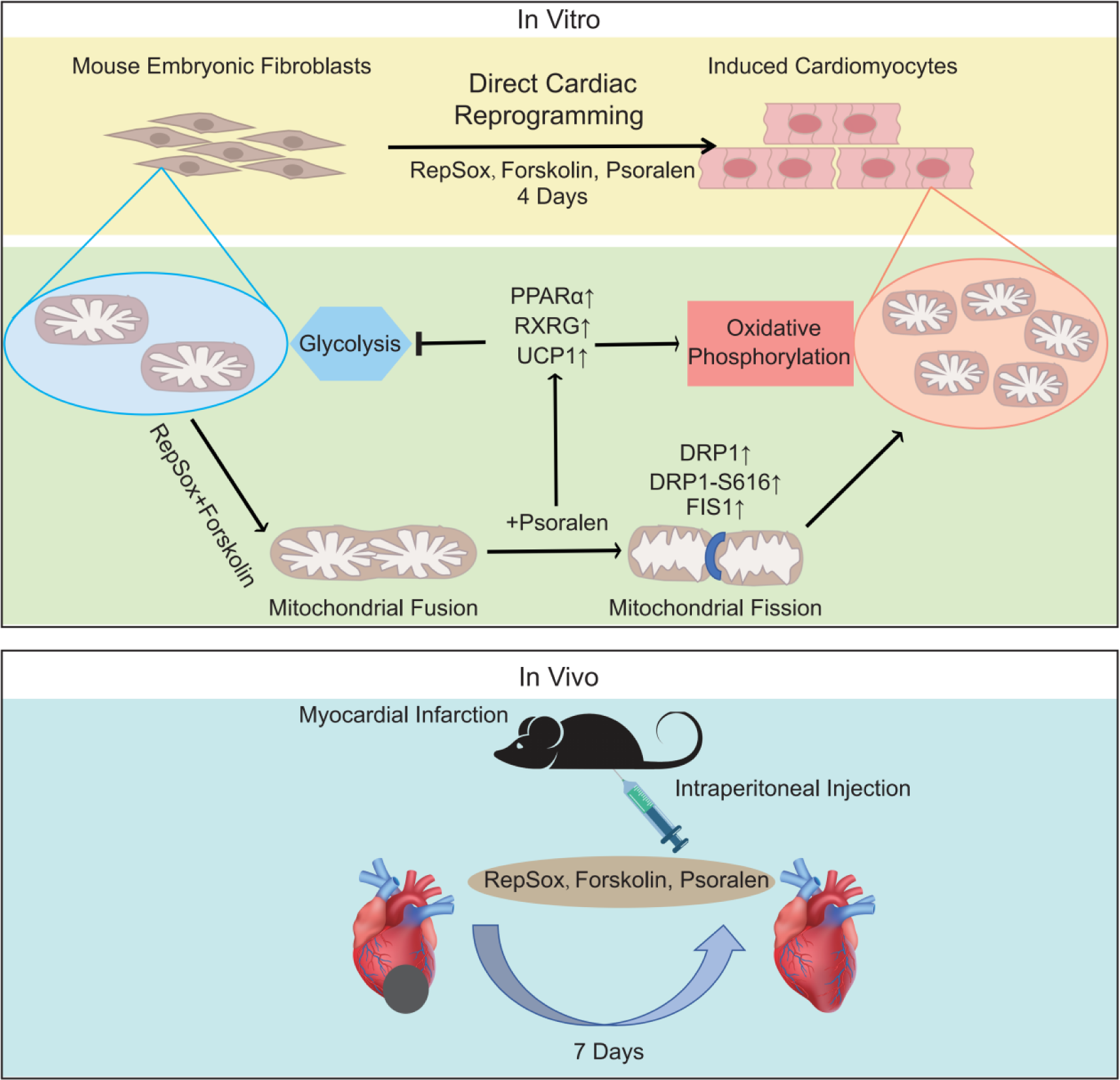

## 1. INTRODUCTION

In recent years, direct cardiac reprogramming of fibroblasts into cardiomyocytes has emerged as a promising strategy for heart regeneration. Chemically-induced direct reprogramming holds great therapeutic potential as it avoids the use of exogenous genetic material and offers the advantage of standardized production^[1]^.

Current approaches primarily utilize cocktails of small molecules that modulate key developmental pathways, including transforming growth factor-β (TGF-β), Wnt, BMP, and fibroblast growth factor (FGF) signaling, to generate induced cardiomyocytes (iCMs)^[2, 3]^. However, a major limitation of these protocols is their requirement for as many as 6-8 molecules (e.g., the CRFVPTZ cocktail comprising CHIR99021, RepSox, Forskolin, VPA, Parnate, TTNPB, and DZnep) and timelines extending to 24 days^[4]^. Thus, developing more robust and efficient direct cardiac differentiation chemical cocktails is crucial for advancing the therapeutic application as well as improving our understanding of key regulatory machineries of heart development.

A critical bottleneck to achieving functional iCM maturation is the metabolic and bioenergetic state of the reprogrammed cell recapitulation of the adult metabolic state^[5]^. Much work has investigated the genetic,epigenetic,and, more recently, translational mechanisms of cardiac reprogramming^[6]^. However, little is known about mitochondrial or metabolic barriers to iCM generation. While mature cardiomyocytes rely predominantly on mitochondrial oxidative phosphorylation (OXPHOS) for energy production, fibroblasts depend on glycolysis^[7]^. Successful direct cardiac reprogramming must involve a bioenergetic switch from glycolysis to OXPHOS, a process intrinsically linked to extensive mitochondrial remodeling^[8]^. Mitochondria are a dynamic organelle that alter their morphology by undergoing fusion and fission processes, maintaining the stability of cell physiological functions^[9]^. Mitochondrial fission, coupled with PINK1-Mfn2-Parkin-mediated mitophagy, is a beneficial and essential physiological process during the perinatal cardiac developmental window. This mechanism directs the removal of fetal mitochondria, enabling their replacement with mature organelles and ensuring the metabolic transition from carbohydrate to fatty acid metabolism critical for postnatal survival.^[10]^ In mouse cardiac fibroblasts, knockdown of the mitochondrial fission-promoting factor mitochondrial fission regulator1-like protein (Mtfr1l) facilitated reprogramming to iCMs by enhancing mitochondrial fusion, suggesting that modulating mitochondrial dynamics can directly influence cell fate conversion^[11]^.

We therefore reasoned that a compound capable of tuning mitochondrial dynamics could address this metabolic bottleneck and enhance iCMs maturation. Based on this premise, we investigated Psoralen (Psr), the main active molecule and quality control component of the traditional Chinese herb medicine Fructus Psoraleae which was registered as a health-care food by the ministry of Public Health of China^[12]^. Emerging evidence suggests that Psr can intervene in the mitochondrial fusion-fission balance and is closely associated with cellular energy metabolism^[13]^. We thus hypothesized that Psoralen enhances direct cardiac reprogramming by modulating mitochondrial dynamics to facilitate the essential metabolic switch.

Psoralen exhibits antioxidant properties and modulates signaling pathways including cAMP^[14]^ and TGF-β^[15]^, which are critical for cell differentiation and metabolism^[16]^. Preliminary work from our group revealed that supplementation of psoralen significantly enhances the efficiency of the classical Forskolin plus RepSox (RF) chemical cocktail in reprogramming mouse embryonic fibroblasts (MEFs) into iCMs, as evidenced by increased expression of cardiomyocyte markers, early emergence of spontaneously beating cells, and improved sarcomeric organization. Despite these encouraging findings, the underlying mechanisms by which psoralen promotes cardiac reprogramming remain largely unclear. Our recent transcriptomic analyses indicate that psoralen treatment upregulates genes associated with fatty acid metabolism, sarcomere assembly, and mitochondrial function, with PPARα, RXRG, and UCP1 identified as key regulatory nodes. These data suggest that psoralen may enhance cardiac reprogramming through coordinated metabolic remodeling and modulation of mitochondrial dynamics, creating a permissive environment for fibroblast-to-cardiomyocyte conversion.

Here, we aimed to systematically investigate the role of psoralen in enhancing chemical cardiac reprogramming. Specifically, we sought to optimize the RF+psoralen combination, assess its effects on iCM structural and functional maturation, and explore the underlying metabolic and mitochondrial mechanisms. Understanding these processes will not only provide mechanistic insights into direct cardiac reprogramming but also offer a potential small molecule-based strategy for heart regeneration.

## 2. MATERIALS AND METHODS

### 2.1 Reagents

The materials used included: DMEM/F12 (1:1) basal medium (1X) (Gibco, 6123073), DMEM basic medium (1X) (Gibco, 6125275), fetal bovine serum (FBS, Procell, 164210-50), MEM non-essential amino acid (Servicebio, G4219), Sodium L-Ascorbyl-2-Phosphate (OriLeaf, S67343), Insulin-Transferrin-Selenium (ITS) G supplement (Servicebio, G4028), Cardiac Troponin T Polyclonal antibody (Protientech, 15513-1-AP), Alpha Actin Polyclonal antibody (Protientech, 23660-1-AP), Beta-actin polyclonal antibody (Protientech, 20536-1-AP), PPARA Monoclonal antibody (Protientech, 66826-1-Ig), RXRG Polyclonal antibody (Protientech, 11129-1-AP), UCP1 Polyclonal antibody (Protientech, 23673-1-AP), DRP1 (C-terminal) Polyclonal antibody (Protientech, 12957-1-AP), FIS1 Polyclonal antibody (Protientech, 10956-1-AP), DRP1 (phospho S616) Monoclonal antibody [EPR27387-57] (abcam, ab314755), Alexa Fluor 594-labeled goat anti-rabbit IgG (Servicebio, GB28301), Alexa Fluor 488-labeled goat anti-rabbit IgG (Servicebio, GB25303), Alexa Fluor 488-labeled goat anti-mouse IgG (Servicebio, GB25301), DAPI staining reagent (Servicebio, G1012), Fluo-4 AM (ThermoFisher, F14201), Mito-Tracker Red CMXRos (Beyotime, C1035), Hoechst 33342 (Solarbio, B8040), First Strand cDNA Synthesis Kit (RNase H minus, Beyotime, D7168M), RepSox (GLPBIO, GC16793), Forskolin (GLPBIO, GC11920), Psr (Beijing Bethealth People Biomedical Technology Co.,Ltd, 23010802), One-Step Western Kit HRP (CWBio, CW2029M), high-strength RIPA buffer (Solarbio, R0010-20ml), BCA Protein Assay Kit (Solarbio, PC0020), SuperSignal™ West Pico PLUS Chemiluminescent Substrate (Thermo Fisher Scientific, 34577), and qPCR SYBR Green Master Mix (High Rox Plus, Yeasen, 11203ES08). Primers were synthesized by Sangon Biotech.

### 2.2 Extraction and Primary Culture of MEFs

MEFs were isolated from embryonic day 13.5 (E13.5) C57BL/6J mouse fetuses as described. Briefly, pregnant dams (10 weeks old, 18-20 g) were euthanized under anesthesia.The uteri were surgically excised under sterile conditions, and fetuses were isolated. After removing fetal heads, limbs, and internal organs, the remaining trunks were rinsed with PBS and minced into fragments (<1 mm^3^). Tissue digestion was performed using 0.25% trypsin at 37°C for 15 minutes. The resultant cell suspension was centrifuged and resuspended in DMEM/F12 medium supplemented with 10% FBS and 1% penicillin/streptomycin, then cultured in T25 flasks. Cells at passage 4 (P4) were utilized for subsequent experiments.

### 2.3 Cell Grouping and Treatment

Passage-4 MEFs were seeded in 12-well plates at a density of 1×10^5^ cells per well. Upon reaching approximately 80% confluency, cells were treated with Cardiac induction medium (CM), consisting of DMEM basic medium supplemented with 10% FBS, 1% penicillin/streptomycin, 50mg/L Sodium L-Ascorbyl-2-Phosphate, 1%ITS and 1%MEM non-essential amino acid. Different small molecules, including Forskolin (F, 10 μM), RepSox (R, 10 μM), and varying concentrations of Psr (5–50 μM), were added to the CM as needed. The culture medium was refreshed every 2 days.

### 2.4 Reverse Transcription Quantitative PCR

Total RNA was extracted using TRIzol reagent from cells. Subsequently, 500 ng RNA was reverse-transcribed into cDNA using the First Strand cDNA Synthesis Kit. Gene expression analysis was performed by RT-qPCR using SYBR Green Master Mix on a CFX96 Real-Time PCR Detection System (Bio-Rad). Reactions were performed in triplicate under the following cycling parameters: initial denaturation at 95 °C for 30 s, then 40 cycles comprising denaturation at 95 °C for 10 s, annealing/extension at 60 °C for 30 s, and melting curve analysis. Expression levels of Ryr2, Tnnt2, β-MHC, Myh6, Myl7, Nkx2.5, Opa1, Mfn1, Mfn2, Drp1, Fis1 and Gapdh were analyzed. Primer sequences are listed in Supplement Table 1.

### 2.5 Immunofluorescence

Cells were fixed with 4% PFA for 10 minutes, washed with PBS, and permeabilized using 0.5% Triton X-100 for 10 minutes. Following blocking with 5% BSA for 1 h, cells were incubated overnight at 4 °C with primary antibodies (cTnT, 1:400; cTnI, 1:200; α-Actin, 1:400) diluted in 5% BSA. Alexa Fluor 594-conjugated goat anti-rabbit IgG (1:400), Alexa Fluor 488-conjugated goat anti-rabbit IgG (1:400) or Alexa Fluor 488-conjugated goat anti-Rabbit IgG (1:400) were applied at room temperature for 1 h. Nuclei were counterstained with DAPI. Fluorescence images were obtained using a Confocal laser scanning microscope (Olympus FV3000), and mean fluorescence intensity was quantified with ImageJ software.

To further investigate mitochondrial function, we quantified mitochondrial mass, as measured by MitoTracker staining.On day 4, the cells were incubated with 100 nM MitoTracker Red CMXros, and 1 μg/mL Hoechst for 20 min at room temperature and then imaged using confocal microscopy.

### 2.6 RNA-sequencing

Total RNA was extracted from cell by Trizol reagent (Invitrogen) separately. The RNA quality was checked by Bioanalyzer 2200 (Aligent) and kept at -80℃.The RNA with RIN >6.0 is right for rRNA depletion. The cDNA libraries were constructed for each pooled RNA sample using the NEBNext® Ultra™ Directional RNA Library Prep Kit for BGI according to the manufacturer’s instructions. Generally, the protocol consists of the following steps: depletion of rRNA and fragmented into 150-200 bp using divalent cations at 94 ℃ for 8 min. The cleaved RNA fragments were reverse-transcribed into first-strand cDNA, second-strand cDNA synthesis, fragments were end repaired, A-tailed and ligated with indexed adapters. Target bands were harvested through AMPure XP Beads(beckman coulter). The products were purified and enriched by PCR to create the final cDNA libraries and quantified by Agilent2200. The tagged cDNA libraries were pooled in equal ratio and used for 150 bp paired-end sequencing in a single lane of the BGI T7. We applied EBSeq algorithm to filter the differentially expressed genes, after the significant analysis, Pvalue and FDR analysis under the following criteria^[37]^.

### 2.7 Western blot (WB) Analysis

Cells were lysed on ice with RIPA buffer containing protease and phosphatase inhibitors, followed by centrifugation at 12,000 g for 10 minutes at 4°C. Protein concentration was quantified using the BCA Protein Assay Kit.Equal amounts of protein (30 μg) were separated by 10% SDS-polyacrylamide gels, transferred onto PVDF membranes, and blocked for 5 minutes. Membranes were incubated with primary antibodies (PPARα, 1:100; RXRG, 1:1000; UCP1, 1:1000; β-actin, 1:2000;DRP1(phospho S616), 1:100; DRP1, 1:1000; FIS1, 1:1000) for 45 minutes at room temperature. After washing, HRP-conjugated secondary antibodies (provided in the One-Step Western Kit) were applied. Protein bands were visualized using SuperSignal™ West Pico PLUS Chemiluminescent Substrate and imaged using the ChemiDOCTM XRS+ Imaging System (Bio-Rad).

### 2.8 Myocardial Ischemia Surgery and treatment

The mice model of Myocardial Ischemia was performed as described previously. Male C57BL/6 mice (8-12 weeks old) were anesthetized by 1% sodium pentobarbital inhalation and ventilated mechanically. After left thoracotomy, the LAD was reversibly ligated with a slipknot (6-0 silk sutures). Sham-operated mice subjected to equivalent procedure with the absence of myocardial ischemia. Mice were randomized to 6 groups: sham group, Model+1% DMSO group, Model+Psr group, Model+RF group, Model+RF+Psr group and Model+Metoprolol group.

To determine anti-fibrotic remodeling of RF+Psr, mice were consecutively treated by intraperitoneal injection or intragastric administration for 7 days post reperfusion injury.

### 2.9 Measurement of Cytosolic Ca2+ Transients

Isolated cardiomyocytes were loaded with 2 μM Fura-4 AM (Sigma-Aldrich) in the dark for 30 min at 22 ± 2°C. Cells were washed, resuspended twice in PBS and placed in a cell chamber. Myocytes were stimulated to contract at a pacing frequency of 1 Hz with 4 ms of electrical stimulation. Myocytes were exposed to 340 or 380 nm excitation wavelengths, and the emitted fluorescent signal was detected at 510 nm. Sarcomere length and fluorescence intensity (a proxy of Ca2+ concentration) were synchronously recorded with a cell contraction-ion detection system (IonOptix, Westwood, MA, United States). Contractility parameters including amplitude, peak time, systolic half-time of decay (T50), diastolic T50, and myofilament sensitivity were measured. Ca2+ transient parameters including amplitude, maximum ascending and descending velocity, and Ca2+ decline time constant were also recorded.

### 2.10 Echocardiography

Mouse echocardiography was performed as previously described. Briefly, hair removal cream was used to expose the anterior chest wall and mice were anaesthetized with 5% isoflurane induction and 0.5% isoflurane maintenance. Transthoracic echocardiography (M-mode and two-dimensional) was performed using a Vevo 2100 (VisualSonics Inc.) high-frequency ultrasound instrument with simultaneous electrocardiogram acquisition via small-needle electrodes. The heart rate was maintained above 500 beats per minute. Researchers blinded to mouse genotypes measured the heart rate, left ventricular internal dimension during diastole and systole (LVIDd/s), end-diastolic interventricular septal thickness in diastole and left ventricle posterior wall thickness in diastole from echocardiographic recordings. Left ventricular fractional shortening percentage, a measure of cardiac contractility, was calculated as the average of three (LVIDd−LVIDs)/LVIDd measurements per animal; n = 6 mice per group.

### 2.11 Histologic Examination

Hearts were harvested and washed in phosphate buffer solution, fixed in 4% paraformaldehyde overnight, and embedded in paraffin. Each paraffin-embedded heart was cut into 4-µm thick sections through the infarction area and stained with hematoxylin and eosin (H&E) for morphological observation. Specimens were stained with Masson’s trichrome stain to evaluate collagen volume. Sections were imaged using a stereomicroscope (Olympus SZ61, Tokyo, Japan).

### 2.12 Statistical Analysis

Data are presented as mean ± standard deviation (SD). Statistical analyses were performed using GraphPad Prism 7.0. All datasets were assessed for normality distribution using Shapiro-Wilk tests (for n < 50), and homogeneity of variance was verified using Brown-Forsythe (for ANOVA). Data comparisons were performed using the statistical tests (Student’s t-test, one-way ANOVA, or two-way ANOVA with appropriate post-hoc tests). Sample sizes (n) for each statistical test are noted in the corresponding figure legends. Statistical significance was set at a threshold of p < 0.05.

## 3. RESULTS

### 3.1 Psoralen Synergizes with the RF Cocktail to Potentiate Cardiac Reprogramming

To determine the optimal concentration of Psoralen (Psr), we first treated mouse embryonic fibroblasts (MEFs) with the RF cocktail supplemented with varying concentrations of Psr (5, 10, 25, and 50 μM). After 4 days, we quantified the expression of key cardiomyocyte maturation markers Ryr2 and Tnnt2 mRNA. Treatment with 10 μM Psr (RF+Psr) yielded the highest mRNA expression levels of both genes (Figure 1A), establishing this as the optimal concentration for all subsequent experiments. Strikingly, spontaneously beating cells or clusters were observed as early as 24 hours after induction with the optimized RF+Psr cocktail—markedly earlier than the previously reported earliest appearance of beating cells (days 6–8) ^[10]^, indicating a substantial acceleration of the reprogramming process.

**Figure 1.**
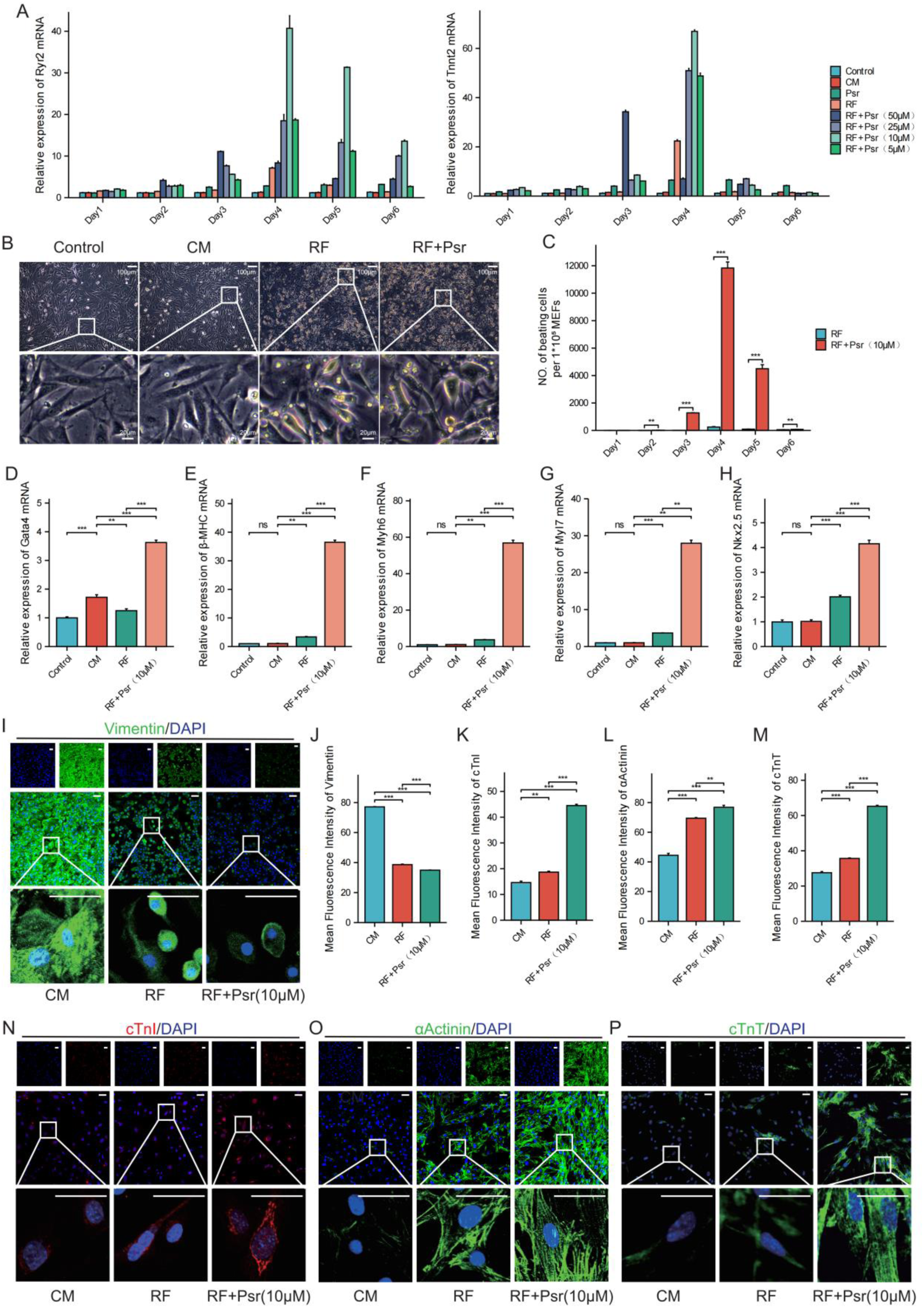
Psoralen(Psr) synergizing with RepSox and Forskolin enhances direct cardiac reprogramming in mouse embryonic fibroblasts. A, Reverse transcription quantitative polymerase chain reaction (RT-qPCR) analysis of Ryr2 and Tnnt2 expression in mouse embryonic fibroblasts (MEFs) cultured in control medium (control) cardiomyocyte differentiation medium (CM), or treated with RepSox + Forskolin (RF) or RF + Psoralen (RF+Psr) from day 1 to day 6 (n=3 per group). B, Representative bright-field images showing morphological changes of cells on day 4 of direct cardiac reprogramming. C, Quantification of spontaneously beating cells from day 1 to day 6 in RF- and RF+Psr-treated groups (n = 3 per group) D-H, mRNA expression levels of Gata4, β-MHC, Myh6, Myl7, and Nkx2.5 in MEFs under the indicated conditions (n = 3 per group). I-P, Immunofluorescence staining images and quantification of Vimentin, cardiac troponin I (cTnI), α-actinin, and cardiac troponin T (cTnT) in MEFs cultured in CM or treated with RF or RF+Psr. Scale bars, 50 μm (n=3 per group).All data are presented as mean ± SEM. *P<0.05, **P<0.01, ***P<0.001.

On day 4, Control and CM-cultured cells retained a spindle-shaped fibroblast morphology with weak refractility. In contrast, cells treated with RF or RF+Psr adopted an enlarged, flattened morphology with strong refractility and reduced nuclear size (Figure 1B). The number of spontaneously beating cells was substantially higher in the RF+Psr group (∼1.2 × 10⁴ per 10⁵ cells) than in the RF group (∼2.5 × 10³ per 10⁵ cells) (Figure 1C and Video S1).

We further validated the increased expression of cardiomyocyte differentiation and functional maturation markers by RT-qPCR (Figure 1D-1H). CM culture significantly upregulated only Gata4 (p<0.001 vs. Control). In contrast, RF treatment markedly enhanced the expression of all five markers—Gata4, β-MHC, Myh6, Myl7, and Nkx2.5—compared to the CM group. Critically, RF+Psr combination therapy further significantly boosted the expression of all markers versus RF alone (p<0.01 for Myl7; p<0.001 for all others).

This molecular progression was confirmed at the protein level by immunofluorescence (IF) staining, which illustrated the loss of fibroblast identity and acquisition of a cardiomyocyte phenotype. Staining for the fibroblast marker Vimentin revealed high expression in the CM group, which was reduced in the RF group and was further diminished to merely ∼32% of the initial level in the RF+Psr group, as quantified by mean fluorescence intensity (MFI; Figure 1I, 1J). Conversely, staining for cardiomyocyte-specific markers showed a corresponding increase. RF treatment alone substantially upregulated cardiac troponin I (cTnI), alpha-actinin, and cardiac troponin T (cTnT) compared to the CM group. The RF+Psr combination induced the most significant upregulation of all three markers (cTnI MFI ∼45; alpha-actinin MFI ∼75; cTnT MFI ∼65), significantly outperforming the RF group (Figure 1K–M). Furthermore, a notable subset of RF+Psr-treated cells exhibited clearly visible sarcomere structures under high-magnification IF imaging, a feature that was rarely observed in cells treated with RF alone (Figure 1N–P).

Taken together, these data demonstrate that Psr (10 μM) significantly enhances cardiac reprogramming by coordinately activating cardiac gene expression and facilitating the assembly of mature contractile units as early as day 4..

### 3.2 Psr Enhances Ultrastructural and Functional Maturation of iCMs

The enhanced reprogramming efficiency induced by Psr was accompanied by improved structural and functional maturation of iCMs.

Transmission electron microscopy (TEM) revealed pronounced ultrastructural differences (Figure 2A). Representative images revealed that cells in RF+Psr group acquired well-organized sarcomeric structures, characterized by aligned myofibrils and distinct Z-discs, which were absent in the CM and RF groups.. Furthermore, mitochondria in RF+Psr treated cells were more abundant and frequently exhibited dumbbell-shaped morphology, a hallmark of active mitochondrial fission..This striking morphological shift, from the sparse and elongated mitochondria observed in RF-only cells to the fission-active state in RF+Psr cells, suggests that Psr plays a critical role in enhancing mitochondrial dynamics during the direct cardiac reprogramming.

**Figure 2.**
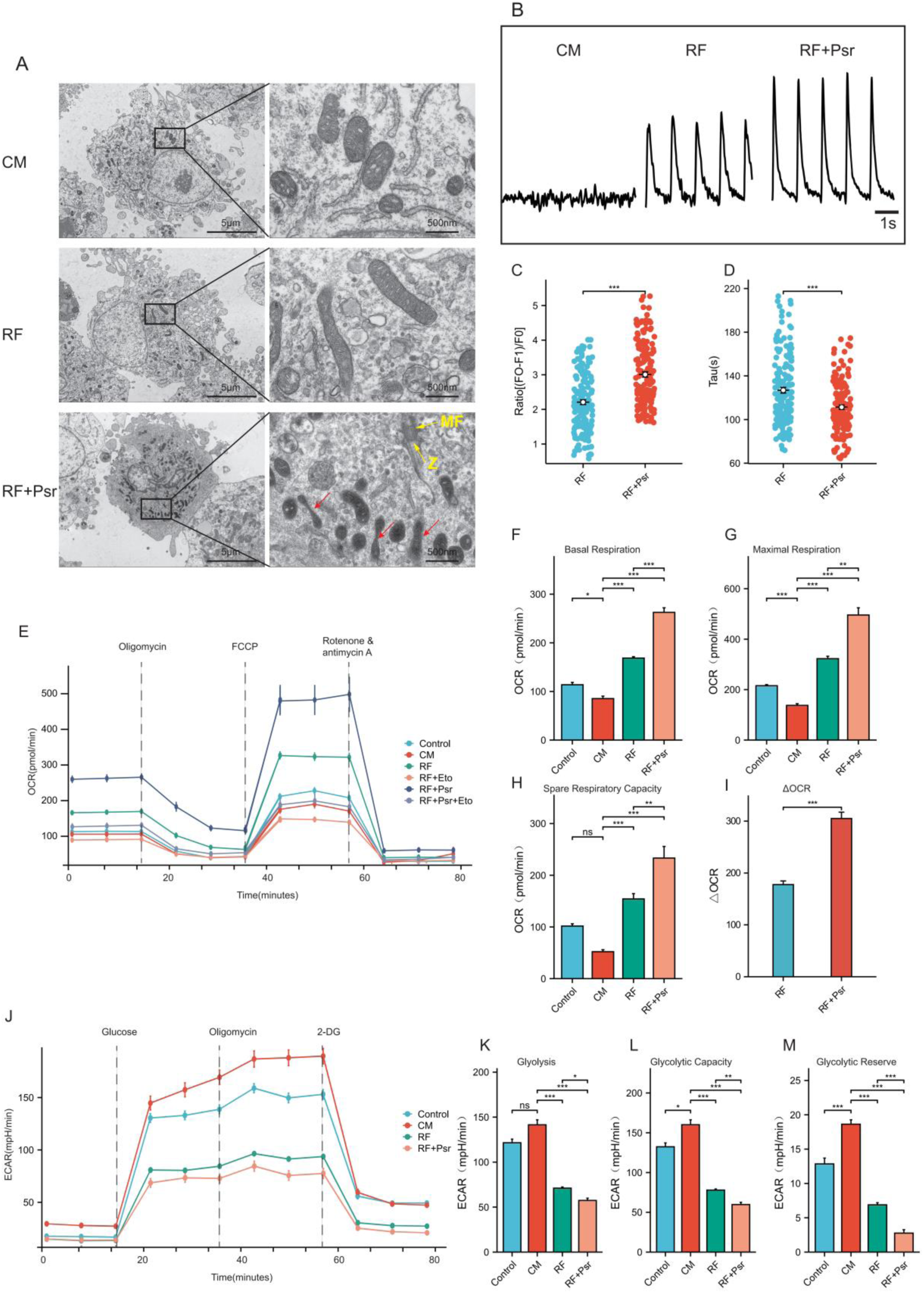
Psr (10μM) leads to functional improvement of iCMs. A, Representative transmission electron microscopy (TEM) images of MEFs cultured in CM, or iCMs generated with RF or RF+Psr. Yellow arrows indicate myofibrils and Z-discs; red arrows indicate dumbbell-shaped mitochondria. B, Representative calcium transient traces recorded from Fluo-4-loaded cells in the indicated groups. C-D, Quantitation of measured Ratio [(F1-F0)/F0] and decay time constant in iCMs from the RF (n = 161) and RF+Psr (n = 148) groups. E, Representative oxygen consumption rate profile analyzed from mito-stress Seahorse assay in the indicated groups (n=5 per group). F-I, Quantification of basal respiration, maximal respiration, spare respiratory capacity in the indicated groups (n=5 per group)and △OCR in RF and RF+Psr treated group. J, Representative extracellular acidification rate profile analyzed from glucose-stress Seahorse assay in the indicated groups (n=5 per group). K-M, Quantification of glycolysis, glycolytic capacity, and glycolytic reserve in MEFs, MEFs cultured in CM, RF and RF+Psr group (n=5 per group). All data are expressed as mean ± SEM. *P<0.05, **P<0.01, ***P<0.001. 2-DG indicates 2-deoxy-glucose; ECAR, extracellular acidification rate; FCCP, fluoro-carbonyl cyanide phenylhydrazone; Eto, Etomoxir; and OCR, oxygen consumption rate; Red arrowheads, Dumbbell-shape mitochondria; Z, Z-dics; MF, myofibrils.

These ultrastructural changes correlated with enhanced calcium handling capacity. The beating iCMs of RF and RF+Psr group exhibited calcium transients (Figure 2B). High-speed video acquisition of Fluo-4-stained cells revealed that iCMs under RF+Psr treatment markedly enhanced these transients, resulting in a substantially higher peak amplitude(Figure 2C) along with a shorter decay time constant (tau) (Figure 2D) compared to RF iCMs. These findings demonstrate Psr enhances calcium cycling during cardiac direct reprogramming process.

To examine the metabolism phenotypes associated with mitochondrial changes, we performed Seahorse metabolic flux assays of MEFs, MEFs cultured in CM with or without RF and RF+Psr(Figure 2E). The oxygen consumption rate measurements showed substantially increased basal, maximal respiration rate, spare respiratory capacity, and change in oxygen consumption rate in iCMs with RF+Psr (Figure 2F-2I). Increased mitochondrial consumption could be a result of both increased mitochondrial number and increased mitochondrial activity(Figure 2J). Moreover, we measured extracellular acidification rate and observed lower glycolysis, glycolytic capacity, and glycolytic reserve in iCMs after RF+Psr treatment (Figure 2K-2M). Reduced glycolysis and increased oxygen consumption in RF+Psr iCMs indicated that Psr may promote the metabolic switch into adult cardiomyocyte-like mitochondrial respiration.

Taken together, these results demonstrate that the supplementation of Psr significantly enhances the maturation of directly reprogrammed iCMs, both structurally and functionally. This is evidenced by the acquisition of mature ultrastructural features (organized sarcomeres with Z-discs), improved functional calcium handling kinetics (higher peak amplitude and faster decay), and a metabolic shift towards oxidative phosphorylation. Critically, we observed a pronounced increase in mitochondrial fission, indicated by the abundance of dumbbell-shaped mitochondria. We propose that Psr potentiates cardiac reprogramming and maturation primarily by activating mitochondrial fission, which in turn facilitates the metabolic remodeling essential for adult-like cardiomyocyte function.

### 3.3 Transcriptomic Profiling Implicates PPARα Signaling as a Key Mediator of Psr Action

To elucidate the molecular mechanisms underlying psoralen-enhanced cardiac reprogramming, we performed RNA sequencing (RNA-seq). A total of 251 genes were significantly upregulated and 72 genes downregulated upon Psr treatment (adjusted P < 0.01, fold change > 2). Notably, many genes with relatively low expression in RF group were markedly activated in the RF+Psr group (Figure 3B). Gene Ontology analysis for the differentially expressed genes (DEGs) revealed that Psr-upregulated genes were predominantly associated with metabolic processes—including fatty acid metabolism and oxidative phosphorylation—as well as structural components of cardiomyocytes such as myosin complexes and Z-discs (Figure 3C). Gene Set Enrichment Analysis (GSEA) further confirmed enrichment of pathways regulating energy metabolism, with the PPAR signaling pathway, tricarboxylic acid (TCA) cycle, fatty acid metabolism, and pyruvate metabolism among the most significantly activated (Figure 3D)

**Figure 3.**
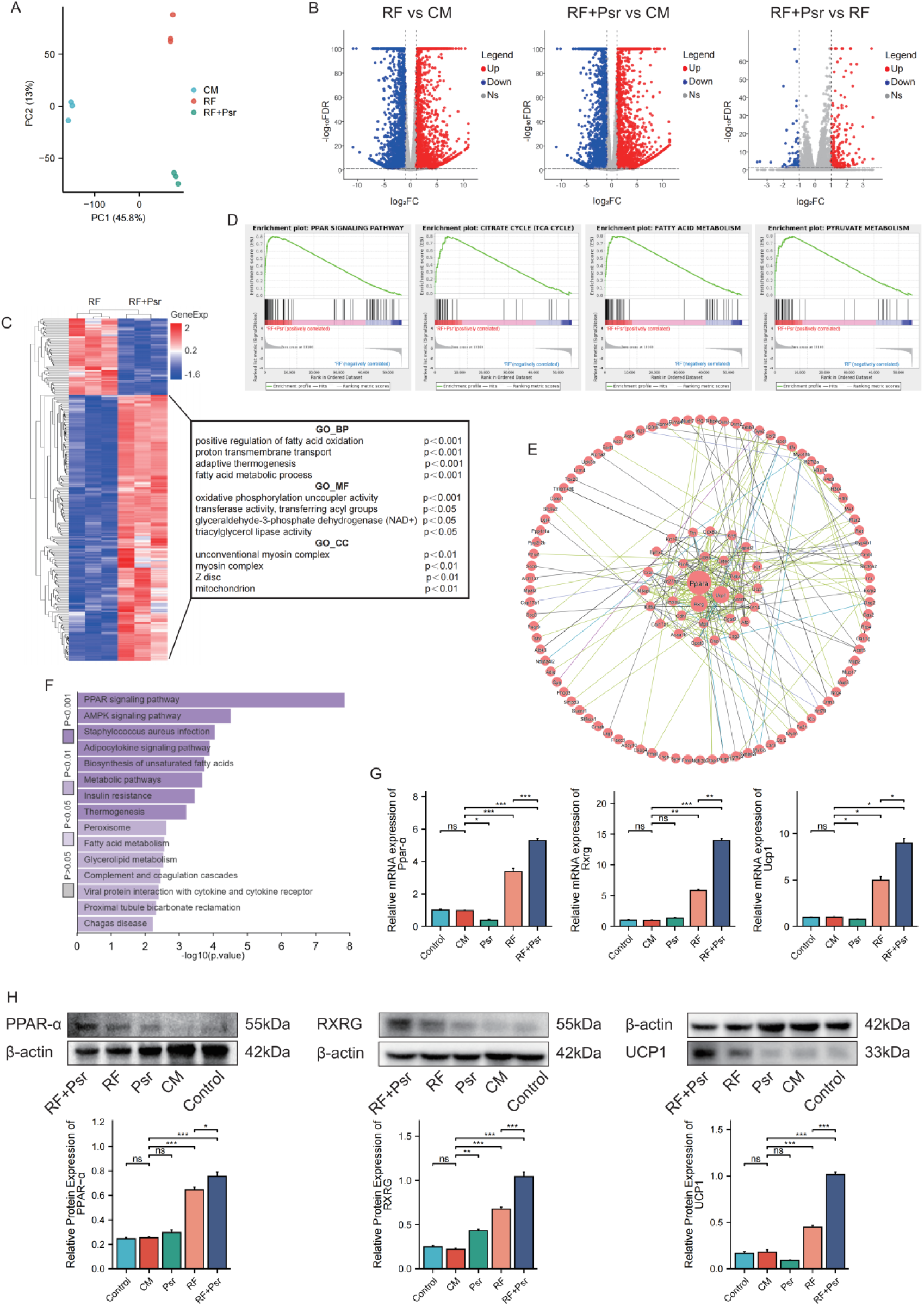
Psr (10μM) activates a PPAR-α/RXRG/UCP1 axis and core metabolic pathways to promote mouse chemical direct cardiac reprogramming. A, Principal components analysis (PCA) of the RNA-Seq data (n=3 per group) from the indicated groups. B, Volcano plot displaying differentially expressed genes (DEGs) between RF+Psr and RF groups.(adjusted P<0.01, fold change >2).C, Heatmap of gene expression for upregulated genes in RF+Psr versus RF. The right-side annotations show representative significantly enriched GO terms for adjacent genes. D, Gene set enrichment analysis (GSEA) of genes upregulated in RF+Psr versus RF. GSEA enrichment plots demonstrate significant activation (NES > 0, FDR < 0.05) of gene sets involved in cellular energy derivation (TCA cycle, pyruvate metabolism) and fatty acid processing (PPAR signaling, fatty acid metabolism). E, PPI network identifies key hub genes. Network of upregulated genes shows Ppar-α, Rxrg and Ucp1 as topologically central hubs, based on their high connectivity, as represented by node size and position. F, Bar plot of pathway enrichment analysis for genes upregulated in RF+Psr versus RF. The bar length represents the degree of enrichment [-log10(P-value)]. Pathways are ranked by their statistical significance (FDR < 0.05). G, RT-qPCR analysis showing mRNA expression of Ppar-α, Rxrg and Ucp1 in Control, CM, Psr, RF and RF+Psr group (n=3 per group). H, Protein expression of key PPAR signaling components under different treatments. Western blotting analysis of PPAR-α, RXRG, and UCP1 protein levels in Control, CM, Psr, RF and RF+Psr group (n=3 per group). β-actin serves as a loading control. All data are expressed as mean ± SEM. *P<0.05, **P<0.01, ***P<0.001.

We next constructed a protein-protein interaction (PPI) network from the upregulated genes to identify key regulatory proteins. The network hubs analysis revealed that PPARα, RXRG, and UCP1 resided at the core of the interactome (Figure 3E). Concurrently, pathway enrichment of the PPI network identified the PPAR signaling pathway as the most significantly enriched (Figure 3F). These results strongly suggest that Psr enhances cardiac reprogramming efficiency, likely through activating the PPAR signaling pathway.

To validate these predictions, we assessed the expression of PPARα, RXRG, and UCP1. Both RT-qPCR and Western blot analysis confirmed that RF+Psr treatment significantly elevated the mRNA (Figure 3G) and protein (Figure 3H) levels of PPARα, RXRG, and UCP1, demonstrating the activation of the PPAR signaling pathway.

Together, these data indicate that Psr supplementation enhances direct cardiac reprogramming efficiency in part via activation of the PPARα/RXRG/UCP1 axis, which may contribute to mitochondrial and structural remodeling observed in iCMs.

### 3.4 Psr Induces Mitochondrial Fission to Facilitate Cardiac Reprogramming

Mitochondria function is critical for cardiomyocyte maturation and is the energetic source for contractility. Given our transcriptomic data indicating enrichment of mitochondrial pathways and activation of the PPAR α/RXRG/UCP1 axis in Psr-treated iCMs (Figure 3), we hypothesized that Psr enhances reprogramming by remodeling mitochondrial dynamics. Supporting this, TEM analysis revealed a high frequency of dumbbell-shaped mitochondria — a hallmark of active fission — in RF+Psr iCMs, contrasting with the elongated and sparse mitochondrial network in RF-only iCMs (Figure 2A).

To quantitatively assess these morphological changes, we performed MitoTracker staining and observed a significant increase in both mitochondrial area and intensity in RF+Psr iCMs (Figure 4A, B). Morphological analysis using ImageJ confirmed that RF+Psr iCMs exhibited a higher proportion of fragmented and donut-shaped mitochondria, while the proportion of elongated (>5 μm) mitochondria was markedly reduced compared with RF-only iCMs. In addition, mitochondrial circularity was increased, and mean branch length was decreased in RF+Psr iCMs, further supporting enhanced fission activity (Figure 4C).

**Figure 4.**
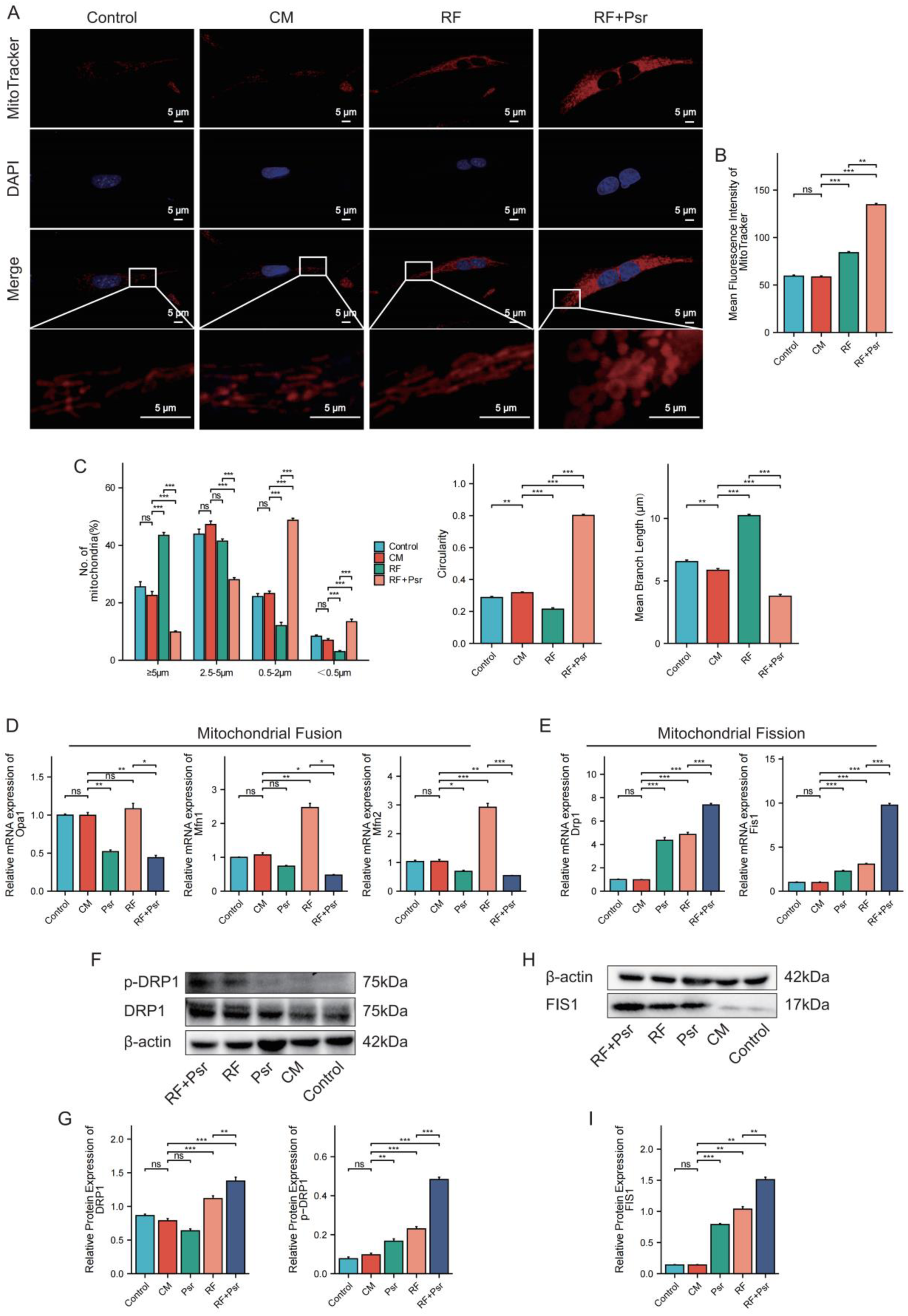
Psr(10μM) enhances mouse chemical direct cardiac reprogramming by promoting mitochondrial fission. A-B, Representative images and quantification of mean fluorescence intensity of mitochondria content labeled using MitoTracker in Control, CM, Psr, RF and RF+Psr group (n=30 per group). C, Quantitative analysis of mitochondrial morphology inControl, CM, RF and RF+Psr group (n=30 per group). D-E, RT-qPCR analysis showing mRNA expression of mitochondrial dynamics regulators Opa1, Mfn1, Mfn2 (fusion) and Drp1, Fis1 (fission) in Control, CM, Psr, RF and RF+Psr group (n=3 per group). F-I, Western blotting of mitochondrial fission markers p-DRP1, DRP1 and FIS1 in Control, CM, Psr, RF and RF+Psr group (n=3 per group). β-Actin was used as the loading control for total protein (DRP1 and FIS1). Total DRP1 protein levels served as the internal control for phospho-DRP1 (p-DRP1). All data are expressed as mean ± SEM. *P<0.05, **P<0.01, ***P<0.001.

RT-qPCR analysis revealed upregulation of fission-related genes (drp1,fis1) accompanied by a reduction in fission-related markers (Opa1, Mfn1, Mfn2) following RF+Psr treatment (Figure 4D, E). Western blotting further confirmed elevated expression of the key fission regulators Drp1 and Fis1 in RF+Psr iCMs relative to the RF group (Figure 4F–I). Collectively, These findings demonstrate that Psr activates a transcriptional and functional program that shifts mitochondrial dynamics toward fission, thereby facilitating the metabolic and structural remodeling essential for cardiac reprogramming.

### 3.5 Synergizing with RF, Psr Attenuates MI-Induced Cardiac Dysfunction and Fibrosis

To evaluate the therapeutic potential of Psr in vivo, we subjected mice to myocardial infarction (MI) and treated them for 7 days. Echocardiographic assessment at day 7 post-MI revealed severe impairment of cardiac function in the Model group, as evidenced by significantly reduced ejection fraction (EF%: Model 38.47±2.78 vs. Sham 71.00±1.78, p<0.001), fractional shortening (FS%: Model 16.55±1.86 vs. Sham 33.52±1.99, p<0.001), left ventricular posterior wall systolic thickness (LVPWs: Model 0.94±0.11 mm vs. Sham 1.81±0.02 mm, p<0.001), and stroke volume (SV: Model 12.81±2.38 μL vs. Sham 36.15±0.47 μL, p<0.001) compared to Sham-operated animals (Figure 5A-E). Notably, treatment with RF+Psr significantly improved all four hemodynamic parameters relative to the Model group (Figure 5B–E), indicating that this combination therapy is more effective than either treatment alone. Among all the therapeutic groups, RF+Psr exhibited the most substantial functional improvement, showing a synergistic effect of Psr in enhancing cardiac function. In contrast, while Psr monotherapy improved FS% (22.97±2.47 vs. Model 16.55±1.86, p<0.01), it did not have a significant impact on the other hemodynamic parameters, emphasizing the importance of combining Psr with RF to achieve a comprehensive therapeutic benefit. Histological analysis further supported the functional improvements, revealing that Metoprolol, RF, and RF+Psr treatments all significantly reduced cardiac injury. This was evidenced by a decrease in infarct size and myocardial fibrosis, as shown by HE and Masson’s trichrome staining (Figure 5F). Among these, RF+Psr treatment provided the most pronounced attenuation of fibrosis and infarct area, consistent with the functional recovery observed in the echocardiographic assessment.

**Figure 5.**
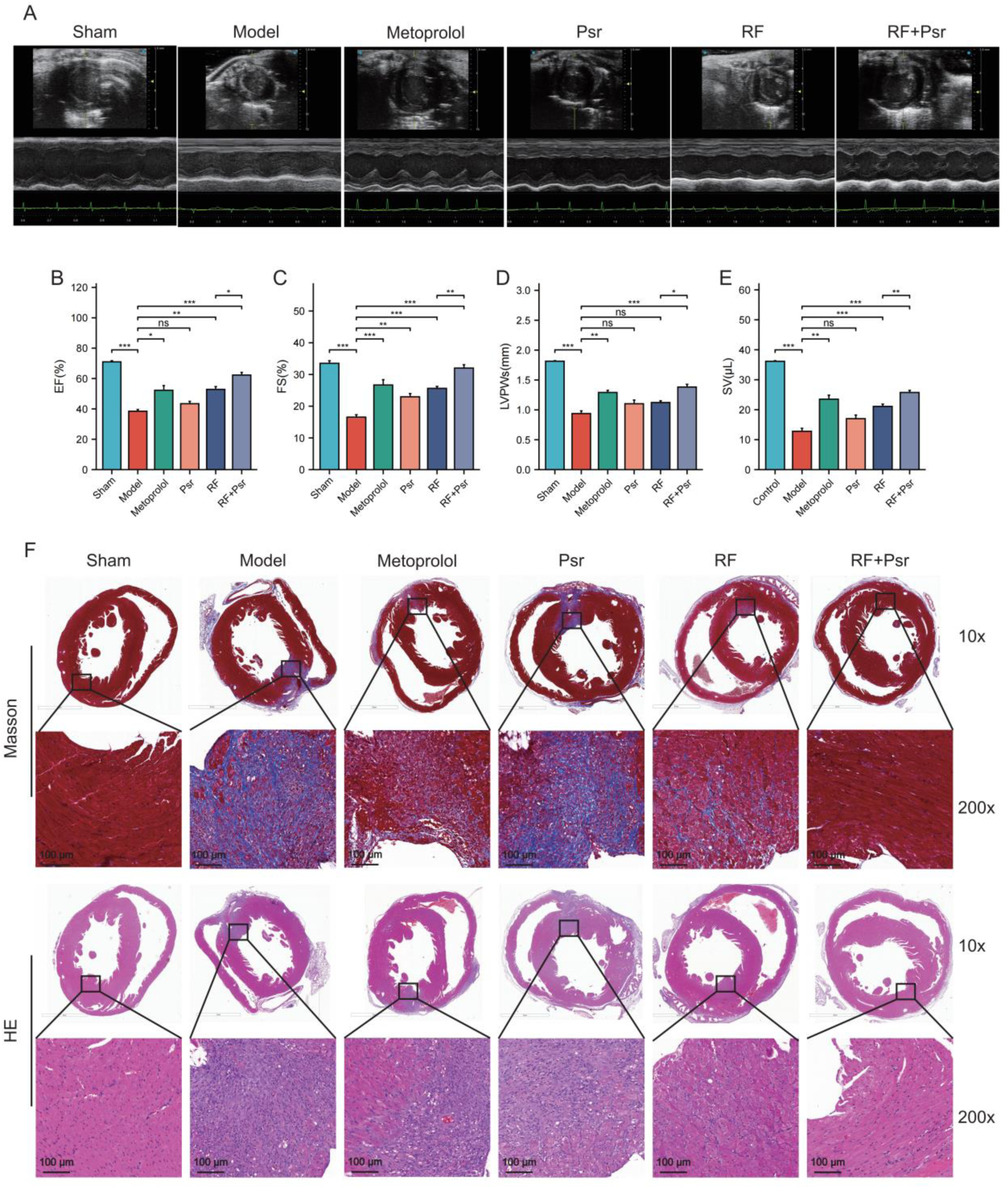
A 7-day treatment with RF+Psr improves cardiac function and ameliorates remodeling after myocardial infarction in mice. A, Representative M-mode echocardiographic images of mice from different groups. B-E, Quantitative analysis of cardiac function parameters: ejection fraction (EF, %), fractional shortening (FS, %), left ventricular posterior wall thickness at end-systole (LVPWs, mm), and stroke volume (SV, μL)(n=6 per group). F, Representative histopathological images of heart sections. Hematoxylin and eosin (HE) staining shows myocardial cell morphology and structure. Masson’s trichrome staining visualizes collagen deposition (blue) and myocardial fibers (red). Sections are from the experimental groups of Sham, Model (Myocardial infarction), Myocardial infarction + Metoprolol (10 mg/kg, p.o.), Myocardial infarction + Psr (10 mg/kg, i.p.), Myocardial infarction + RepSox (10 mg/kg, i.p.)+Forskolin (10 mg/kg, i.p.), Myocardial infarction+ RepSox (10 mg/kg, i.p.)+Forskolin (10 mg/kg, i.p.) + Psr (10 mg/kg, i.p.). All data are expressed as mean ± SEM. *P<0.05, **P<0.01, ***P<0.001.

Together, these findings demonstrate that RF+Psr therapy not only enhances the efficiency of in vitro reprogramming but also offers significant protection against MI-induced cardiac dysfunction and remodeling in vivo. The synergistic effects of Psr with RF underscore its potential as a promising therapeutic approach for heart regeneration.

## 4. DISCUSSION

In this study, we demonstrated that psoralen (Psr), a bioactive component derived from Psoralea corylifolia, significantly enhanced the efficiency and functional maturation of direct cardiac reprogramming induced by RepSox and Forskolin (RF). First, Psr supplementation rapidly generates iCMs within only 4 days, increased the proportion of cells expressing cardiomyocyte markers, accompanied by improved sarcomere organization, calcium transient properties, and spontaneous contractile activity. Second, metabolic assays revealed that RF+Psr-treated induced cardiomyocytes (iCMs) exhibited a pronounced shift from glycolysis toward oxidative phosphorylation (OXPHOS), consistent with a more mature cardiomyocyte metabolic phenotype.Third, RNA sequencing uncovered activation of the peroxisome proliferator-activated receptor (PPAR) signaling pathway, with PPARα, RXRG, and UCP1 identified as critical hub genes. Mechanistically, Psr promoted mitochondrial fission. Finally, the therapeutic potential of this Psr-enhanced reprogramming strategy was confirmed in vivo, where it significantly attenuated MI-induced cardiac dysfunction and fibrosis. Collectively, these findings provide compelling evidence that Psr enhances direct cardiac reprogramming through metabolic reprogramming and mitochondrial remodeling.

Previous studies on direct cardiac reprogramming have primarily focused on transcription factor cocktails, such as Gata4, Mef2c, and Tbx5 (GMT) or their derivatives, which were able to induce partial cardiomyocyte-like phenotypes but with low efficiency and immature functionality^[17]^. More recently, small molecules have been applied to replace or supplement transcription factors, with combinations including CHIR99021, RepSox, Forskolin, and others^[18]^. While these approaches improved feasibility, the efficiency and maturity of iCMs remained suboptimal^[19]^. Our work extends these findings by introducing a natural small molecule, psoralen, as a novel enhancer of chemical reprogramming. Unlike most previously reported small molecules that act through epigenetic modification or signaling modulation, Psr primarily targets cellular metabolism, particularly fatty acid oxidation and mitochondrial function. This mechanism provides a unique and complementary angle compared to earlier strategies.

Our RNA-seq analysis indicates that Psr acts primarily by activating the PPAR signaling pathway, with PPARα, RXRG, and UCP1 identified as critical hub genes. PPARα is a master regulator of fatty acid oxidation and mitochondrial biogenesis^[20, 21]^. Our findings are strongly supported by a recent study demonstrating that enhanced FAO, driven by the PGC1α/PPARβ axis, is a critical determinant for both the efficiency and functional maturation of directly reprogrammed iCMs^[22]^. Moreover, PPARα, in complex with RXRG, acts as a transcriptional regulator of fatty acid oxidation genes, thereby promoting mitochondrial biogenesis and enhancing oxidative ATP production^[23]^. This shift provides a robust energy supply necessary for contractile function and sarcomere assembly. Broader evidence indicates that PPARα activation ameliorates atherosclerotic dyslipidemia and reduces cardiovascular riske^[24]^.

The concomitant upregulation of UCP1, a key mediator of mitochondrial uncoupling^[25]^, provides a plausible link between Psr-induced PPARα activation and the observed mitochondrial remodelin. Specifically, Psr-induced upregulation of UCP1 reduced mitochondrial ROS production, thereby alleviating oxidative stress, which is otherwise a barrier to successful reprogramming^[26]^. Reduced ROS also contributes to genome stability and protein homeostasis, further supporting iCM survival and maturation^[27]^. This connection is strongly supported by findings in brown adipocytes, where UCP1-mediated thermogenesis occurs concurrently with extensive mitochondrial fragmentation driven by Drp1 phosphorylation^[28]^. In line with this, Psr promoted mitochondrial fission. Our observations confirm this, as observed via electron microscopy and validated by MitoTracker assays, which promotes a dynamic mitochondrial network more suited to high oxidative metabolism.

Mitochondria are dynamic organelles that play important roles in various in cell metabolism, proliferation, apoptosis and differentiation^[29]^. Mitochondrial dynamics have been implicated in cell fate regulation, where fission often facilitates reprogramming initiation^[30]^. While excessive fission has been linked to apoptosis, moderate fission may facilitate the redistribution of mitochondria to areas of high energy demand, such as sarcomeres, thereby supporting functional maturation^[31]^. In the context of cardiac reprogramming, studies have observed that successful direct cardiac reprogramming is accompanied by a profound reorganization of the mitochondrial network^[32]^, which involves mitochondrial fission as a fundamental prerequisite for the metabolic reprogramming essential to iCM maturation^[33]^. Mitochondria fission facilitates the selective removal of outdated or dysfunctional mitochondria through mitophagy^[34]^. Previous studies have implicated the importance of mitochondrial fission proteins for the direct cardiac reprogramming of fibroblasts, as the process requires mitophagy activation to clear outdated mitochondria and support the metabolic shift toward oxidative phosphorylation. Consistently, knockdown of essential autophagy genes such as Atg5 significantly impairs iCM generation^[35]^. The critical role of mitochondrial fission extends to heart development and disease in vivo. Mice lacking mitochondrial fission proteins (Drp1 and Mff) exhibit developmental cardiac defects and increased susceptibility to cardiac injury^[36]^, Therefore, our finding that Psr enhances direct cardiac reprogramming by promoting mitochondrial fission is consistent with these established concepts. Moreover, our data confirm this view and further identify UCP1-mediated modulation of mitochondrial homeostasis as an unrecognized mechanism in the context of cardiac reprogramming.

In summary, the mechanistic basis for Psr’s beneficial effects involves coordinated regulation of transcriptional networks, energy metabolism, and organelle dynamics. Integrating these findings, we propose Psr activates the PPARα-RXRG transcriptional complex, leading to enhanced fatty acid oxidation, UCP1-mediated mitochondrial fission, and a metabolic shift toward oxidative phosphorylation. This cascade establishes an optimized energetic and structural foundation that drives the functional maturation of iCMs, including proper sarcomere assembly and contractile capability.

The translational relevance of our findings is underscored by the robust in vivo therapeutic effects. The intraperitoneal administration of the RF+Psr cocktail markedly improved cardiac function and reduced fibrosis after myocardial infarction within just 7 days. The superior performance of the combination therapy over RF or Psr alone highlights a potent synergistic effect and positions Psr as a promising adjuvant for cardiac regeneration strategies.

Despite these advances, our study has limitations. First, the precise molecular target of Psr within the PPAR pathway remains to be fully elucidated. Second, while we established a strong correlation between PPAR activation, mitochondrial fission, and reprogramming success, definitive proof of causality would require loss-of-function experiments to confirm that these nodes are necessary for Psr’s effects. Finally, the efficacy and safety of this approach need to be validated in human cells and in more chronic in vivo models before clinical translation can be considered.

In conclusion, our study unveils Psr as a highly effective molecule for enhancing direct cardiac reprogramming. We delineate a clear mechanistic pathway from PPAR pathway activation to mitochondrial fission and subsequent metabolic and functional maturation of iCMs. This work not only introduces a novel and efficient chemical cocktail for generating iCMs but also fundamentally advances our understanding of how mitochondrial dynamics govern cell fate determination. Targeting mitochondrial fission emerges as a powerful new strategy for heart regeneration.

## Author contributions

Zhiguo Zhang and Ruyuan Zhu acquired the financial support for the project leading to this publication. Ruyuan Zhu conducted research. Wenjie Li carried out the experiments and wrote the article. Xinyu Wan and Ding Cheng carried out the experiments. Zhiguo Zhang, Haixia Liu and Ruyuan Zhu reviewed the manuscript. Wenjie Li is the first authors. Ruyuan Zhu and Zhiguo Zhang contributed equally to this work and should be considered corresponding authors. All authors read and approved the final manuscript.

## Acknowledgments

This study was financially supported by the Youth Talent Promotion Project of Chinese Academy of Traditional Chinese Medicine (grant number: ZZ16-YQ-033), National Natural Science Foundation of China (grant number: 82074297), CACMS Innovation Fund (grant number: CI2021A00107), Basic Scientific Research Fund of Chinese Academy of Traditional Chinese Medicine (grant number: YZX-202410).

## Ethical Compliance Statement

All experimental procedures involving animals were conducted in strict accordance with the Guidelines for the Care and Use of Laboratory Animals and were approved by the Institutional Animal Care and Use Committee of the Laboratory Animal Center, Institute of Basic Theory for Chinese Medicine, China Academy of Chinese Medical Sciences (Approval Certification No. IBTCMCACMS21-2409-06; Acceptance No. IBTCMCACMS-2409006). Every effort was made to minimize animal suffering and to reduce the number of animals used while maintaining scientific validity.

